# In Silico Identification of Aminoadipate Semialdehyde Synthase (*AASS*) as a Novel Prognostic Biomarker in Triple-Negative Breast Cancer

**DOI:** 10.64898/2026.02.19.706804

**Authors:** Mehwish Majeed, Muhammad Zurgham Akram, Hassan Tariq

## Abstract

Triple-negative breast cancer (TNBC) is an aggressive subtype that lacks effective targeted therapies. This study aimed to identify robust prognostic biomarkers by integrating network biology with machine learning (ML) approaches. TNBC expression cohorts were analysed to identify differentially expressed genes (DEGs) and crucial gene clusters via limma and Weighted Gene Co-expression Network Analysis (WGCNA). In results, 579 DEGs were identified, and network analysis revealed two TNBC-associated modules. Overlapping determined 208 genes enriched in cell-cycle and mitotic-regulation pathways. To identify candidate biomarkers, protein-protein interaction (PPI) networks and ML feature selection techniques, including support vector machine with recursive feature elimination module (SVM-RFE) and least absolute shrinkage and selection operator (LASSO) regression, were performed. The Kaplan-Meier (KM) analysis revealed *AASS* and *CCNA2* were favourable prognostic markers, whereas *CXCL8, SPP1*, and *CCNB1* were poor prognostic markers. Multi-level validation and immune-subtype analysis were carried out, revealing *AASS* as a novel TNBC-associated metabolic tumour suppressor.

## 1. Introduction

Breast cancer is a widespread form of carcinoma and poses critical health hazards to women’s health. It is categorised as the subsequent primary reason for death by cancer (Harbeck et al., 2019; Sung et al., 2021). Among its subtypes, triple-negative breast cancer (TNBC) comprises approximately 15 to 20% of all breast cancer diagnoses. Unlike other breast cancer types, TNBC lacks the three main receptor expressions, including estrogen receptor (ER), progesterone receptor (PR), and human epidermal growth factor receptor 2 (HER2) (Garrido-Castro, Lin and Polyak, 2019). Therefore, TNBC has no approved targeted treatment options; only standard chemotherapeutic interventions, surgeries, and radiation therapies remain the accepted standard for treating TNBC patients (Dogra et al., 2025; Yin et al., 2020). This contributes to poor prognosis, metastasis rate, high recurrence, and mortality. To address this, there is an emergent need to investigate novel gene targets. The discovery of effective target genes is crucial for the development of specialised treatments and enhancing the survival rates of TNBC patients.

The evolution of high-throughput sequencing has enabled researchers to analyse changes in gene expression data across multiple cancer types. In medical research, bioinformatics is most commonly used in identifying potential biomarkers in both malignant and non-malignant conditions, aiding clinicians in predicting patient prognosis and treatment response (Brenner, 2019). Weighted gene co-expression network analysis (WGCNA) is a frequently used approach to determine the correlation of gene expression profiles among distinct biological samples (Langfelder and Horvath, 2008), commonly including cancerous and normal tissues. It provides important gene modules and aids in determining hub genes on the basis of the interconnection of gene clusters with clinical traits. Differential Expression Analysis (DEG) uncovers genes that show significant expression differences between conditions, such as disease vs normal samples (Ho et al., 2008).

Machine learning (ML) feature selection methods are fundamental in biomarker identification, as they extract the most significant molecular features from complex, high-dimensional datasets, improving model precision and biological validity (Dhal and Azad, 2021). LASSO regression is among the standard ML methods used for feature extraction and classification of high-dimensional data. Through regularisation, it can efficiently identify important features with non-zero coefficients (Muthukrishnan and Rohini 2016). SVM-RFE is a hybrid feature selection technique (Tariq, Majeed and Ahmad, 2025). Recursive Feature Elimination uses an iterative approach to rank features. In each iteration, it calculates the score of each feature based on specific criteria and removes the feature with the lowest score. This process is repeated until all genes are ranked (Sanz et al., 2018). The identification of hub and key genes in TNBC has been explored in several studies using various bioinformatics approaches. For instance, studies have applied differential expression analysis alongside protein-protein interaction (PPI) networks (Wei et al., 2021; Zhai et al., 2019), while others have integrated differentially expressed genes (DEGs) with KEGG pathway enrichment analysis for hub gene detection and its functional analysis (Chen et al., 2020; Peng et al., 2017). Researchers also explore biomarker discovery by utilising WGCNA combined with Mendelian randomisation (Lin, Wang and Yang, 2024) or WGCNA paired with the LASSO feature selection method (Chen, Cai and Wang, 2022). Furthermore, meta-analysis approaches combined with WGCNA have also been conducted (Cao et al., 2021). Despite these advances in transcriptomic analysis for biomarker discovery in TNBC, ML integration remains underexplored. Integrating transcriptomic network study with differential expression signals and ML-based feature prioritisation would enable the identification of biologically central and novel TNBC biomarkers that may be missed by utilising conventional bioinformatics approaches alone.

To address this gap, this study established an integrative systematic framework that combines WGCNA with DEGs analysis, followed by dual ML feature selection (LASSO and SVM-RFE) and multi-level validation of prioritised candidates. Specifically, we aimed to:

- Identify novel TNBC-associated biomarkers through integration of network topology and differential expression analysis with ML-based feature prioritisation.
- Characterise the biological functions of overlapping genes via enrichment analysis, construct protein-protein interaction networks using STRING, and determine hub genes through topological ranking in Cytoscape (CytoHubba).
- Evaluate prognostic significance of ML-prioritised and network-central genes using survival analysis to assess their clinical relevance.
- Perform comprehensive validation, including functional annotation, immune-context association, and multi-level transcriptomic and proteomic expression analysis of novel candidates.

In conclusion, our research may help discover important genes for TNBC detection and prognosis and develop novel approaches to therapy.

## 2. Materials and methods

### 2.1 Data collection

The gene expression profiles utilised in this study were sourced from the publicly accessible Gene Expression Omnibus (GEO) repository, hosted by the National Centre for Biotechnology Information (NCBI) (http://www.ncbi.nlm.nih.gov/geo). The primary dataset, GSE76250, was generated using the Affymetrix Human Transcriptome Array 2.0 (platform GPL17586) and comprises 165 TNBC samples along with 33 matched normal breast tissue samples. For validation, two additional expression datasets were used: GSE38959, derived from the Agilent GPL4133 Whole Human Genome Microarray 4 x 44K, which contains 30 TNBC and 13 normal samples; and GSE65194, based on the Affymetrix GPL570 platform (HG-U133 Plus 2), from which 55 TNBC and 11 normal breast tissue samples were selected.

### 2.2 Differentially expressed genes screening

The GEOquery package (version 2.72.0) of R was utilised to retrieve the GSE76250 dataset. (version 3.60.6) was used to determine the DEGs between TNBC and normal breast samples (Davis and Meltzer, 2007; Ritchie et al., 2015). A threshold value of adjusted p-value below 0.05 and | log2 FC| ≥ 1 was applied to select DEGs. A volcano plot was plotted using the ggplot2 package (version 3.5.2) to visualise the distribution of differentially expressed genes (DEGs), classified as upregulated, downregulated, or non-significant based on their expression levels (Tollefson, 2021). Further, a heatmap of the top DEGs was generated using the SRplot (Tang et al., 2023), an online visualisation tool. Probe-level annotations were performed using the SOFT annotation file provided with the dataset.

### 2.3 Weighted gene co-expression network construction and determining key modules

We carried out WGCNA on the GSE76250 dataset using the R package “WGCNA” (version 1.73) in order to find co-expressed gene modules and explore their associations with clinical characteristics in TNBC. The expression data were retrieved using the R GEOquery package (Davis and Meltzer, 2007) and processed to remove missing values. Variance analysis was performed to select the top 25% of genes to reduce noise and computational complexity, in which the highest median absolute deviation (MAD) represents the most variable genes across samples. This subset was used to build the co-expression network. The β soft-threshold value was selected by the “pickSoftThreshold” function as it provided a high scale-free topology fit index while maintaining sufficient mean connectivity. Initially, an adjacency matrix was generated to represent gene co-expression relationships. Subsequently, this matrix was converted into a topological overlap matrix (TOM) to measure network interconnectedness. Following this, hierarchical clustering was utilised based on TOM-derived dissimilarity values (1-TOM) to cluster genes into distinct modules, and gene clusters were detected with the help of dynamic tree cutting. Modules having similar expression profiles were subsequently united based on module eigengene correlation. The association between module eigengenes and the phenotypic status (TNBC vs. normal) was evaluated to identify the module of interest. In addition, gene significance (GS) and module membership (MM) were also assessed.

### 2.4 Functional pathway and gene enrichment analysis

GO and KEGG pathway analysis were carried out using ShinyGo (http://bioinformatics.sdstate.edu/go/), an online interactive bioinformatics platform used for functional pathways and gene enrichment analysis (Ge, Jung and Yao, 2019). A total of 208 genes were subjected to analysis, which was obtained by intersecting DEGs against genes from the two WGCNA-identified modules of interest. Enrichment analysis of GO terms was carried out across three categories, which include biological process (BP), molecular function (MF), and cellular component (CC), alongside KEGG pathway enrichment. A threshold of 0.05 for the false discovery rate (FDR) was applied to determine statistically significant results.

### 2.5 Hub gene selection by PPI network mapping

STRING is a publicly available database tool (https://string-db.org/) designed to analyse and predict functionally associated networks among proteins within an organism (Szklarczyk et al., 2022). In this study, PPI networks were generated through this platform based on genes obtained from WGCNA-selected modules intersecting with DEGs. The PPI networks with functional interaction scores ≥ 0.4 were downloaded and visualised with the help of Cytoscape software (https://cytoscape.org/). Subsequently, hub gene identification was performed using the CytoHubba plugin integrated within Cytoscape, applying topological analysis of the PPI networks. The degree score method of CytoHubba was selected, which finds hub genes based on the total direct links associated with each node (Chin et al., 2014; Shannon, 2003).

### 2.6 Identification of key genes through machine learning-based feature selection

Two machine learning-based feature selection methods were employed. First, LASSO logistic regression that selects the key variables by tuning the value of λ that minimises classification error (Muthukrishnan and Rohini, 2016). Second, SVM-RFE, which is based on the SVM framework to identify important features; it iteratively removes less discriminative features and retains only the most significant ones. The R package “glmnet” was used to conduct LASSO regression analysis (Friedman, Hastie and Tibshirani, 2010). Subsequently, SVM-RFE was performed by the “e1071” (version 1.7-16) and “caret” (version 7.0-1) packages. From SVM-RFE, the top ten most informative genes were selected according to their contribution to classification performance. Genes shared between the two methods were considered potential key genes. Receiver Operating Characteristic (ROC) curves and corresponding Area Under the Curve (AUC) metrics for the selected key genes were calculated using the R package “pROC” (version 1.18.5).

### 2.7 Survival prognostic analysis

The KM Plotter (http://www.kmplot.com), a public database that contains patient clinical information and gene expression data, was used to assess the prognostic potential of hub and key genes identified from PPI and machine learning algorithms (Lánczky and Győrffy, 2021). Overall Survival (OS), Distant Metastasis-Free Survival (DMFS), and Relapse-Free Survival (RFS) were evaluated. In the KM plotter, the mRNA breast cancer repository was selected, and then gene-associated probes were chosen following “only JetSet best probe set” to ensure probe quality and consistency. DMFS and RFS patient data were chosen based on ER-/PR-negative status, assessed by immunohistochemistry (IHC), and HER2-negative status by array, and for OS, the St. Gallen basal-like subtype was selected. Patients were split using the platform’s auto-selected best cutoff percentile. Consequently, log-rank p-values under the 0.05 threshold indicated a strong statistical significance.

### 2.8 Cross-dataset validation

Further, two independent datasets, GSE38959 and GSE65194, of TNBC and control samples were analysed to validate our results. On these datasets, DEG analysis was performed using GEO2R with consistent DEG selection criteria (|log2 FC| ≥ 1.0 and adjusted p-value below 0.05). The common DEGs found across all three datasets were represented by Venn diagrams.

### 2.9 Novel candidate validation at the transcriptomic and proteomic level

Tumour suppressor potential of alpha-aminoadipic semialdehyde synthase (*AASS*) in breast invasive carcinoma (BRCA) and its subtype TNBC assessed using TNM-plot (https://tnmplot.com/) and UALCAN (https://ualcan.path.uab.edu) (Bartha and Győrffy, 2021; Chandrashekar et al., 2022). Firstly, in TNMplot analysis, Gene chip non-paired tumour and normal samples, RNA sequencing paired vs adjacent normal tissue samples, BRCA normal vs. tumor expression, BRCA stage-specific analysis, BRCA subtype analysis, and TNBC subtypes analysis were performed. Expression was further analysed across protein-level in BRCA using UALCAN CPTAC (Clinical Proteomic Tumour Analysis Consortium) data. The analysis focused on the major molecular subtypes of BRCA: Luminal A, Luminal B, HER2-enriched, and Basal/TNBC.

### 2.10 Immune and tumour microenvironment analysis

The TISIDB public database (http://cis.hku.hk/TISIDB/) was employed to evaluate *AASS* expression across both immune and molecular subtypes of BRCA (Ru et al., 2019). Statistical significance across groups was measured using the Kruskal-Wallis test, as implemented in TISIDB. The detailed workflow of this research is shown in Figure 1.

**Figure 1.**
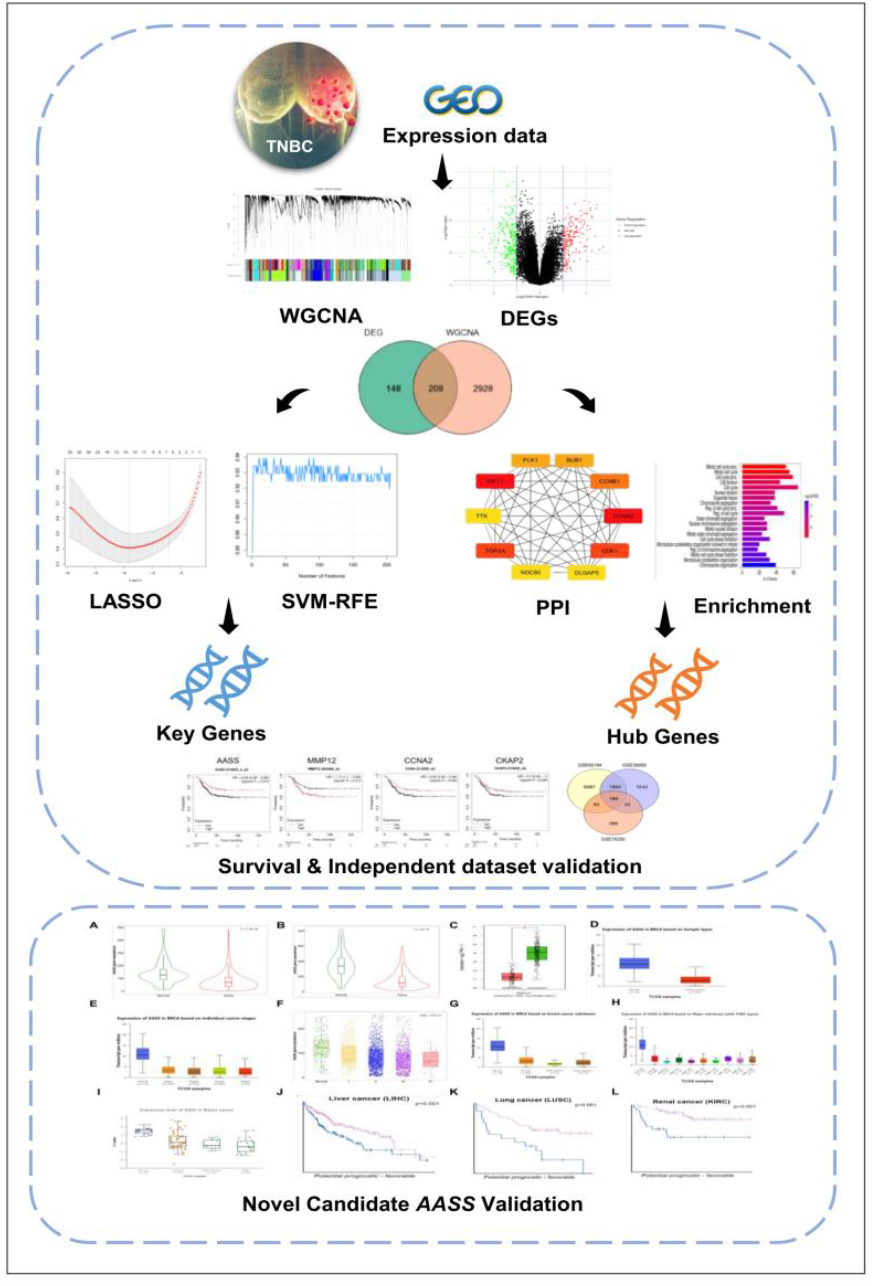
Detailed workflow diagram of the Integrative framework implemented to discover novel biomarkers in TNBC

## 3. Results

### 3.1 Detection of DEGs in TNBC patients

A total of 579 DEGs were recorded from the differential gene expression analysis of TNBC patients. 204 genes showed an upregulated trend, and 375 were downregulated. This indicates significant alterations in gene expression profiles between tumour and normal states. The distribution and significance of these DEGs are visually summarised using a volcano plot in Figure 2(A), which highlights the most prominently altered genes. Additionally, a heatmap presented in Figure 2(B) indicates the expression patterns of these DEGs across all analysed samples.

**Figure 2.**
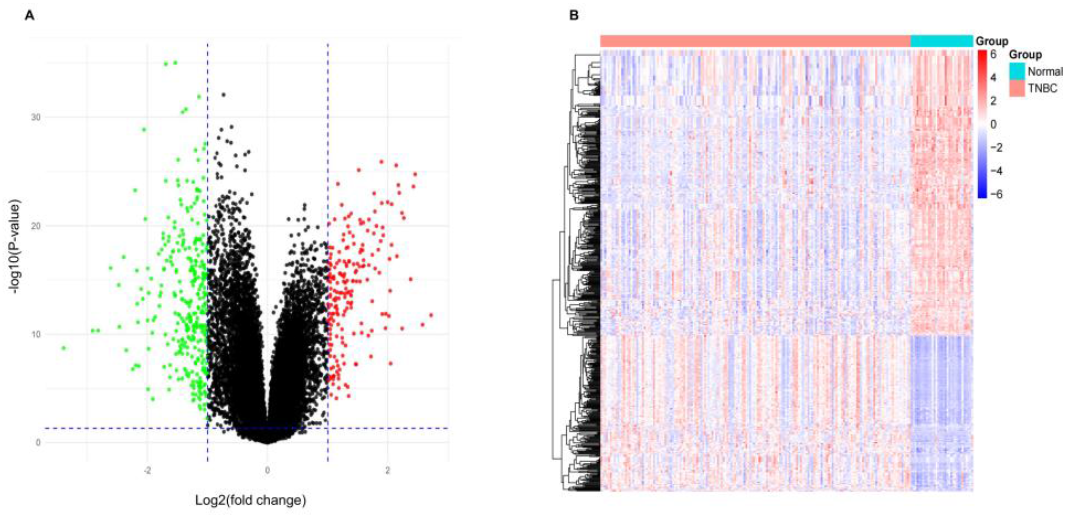
DEGs identification in TNBC patients. (A) volcano plot showing distribution of DEGs, 198 TNBC and normal samples. (B) Heatmap for DEGs, Blue indicates upregulation, red indicates downregulation, and black represents unchanged genes

### 3.2 WGCNA-driven analysis of gene co-expression and module detection

During the clustering step, no outliers were identified. To determine optimal soft-threshold power (β), the network topology was tested across a range of powers from 1 to 20. Using the “pickSoftThreshold” function, a β value of 5 was selected, and it also achieves a scale-free topology fit index above 0.85 while maintaining adequate mean connectivity, shown in Figure 3(A). The β=5 was used to create the adjacency matrix, followed by forming TOM and dis-TOM. In module identification, the dynamic tree cut size was set to a minimum module size of 40 genes.

**Figure 3.**
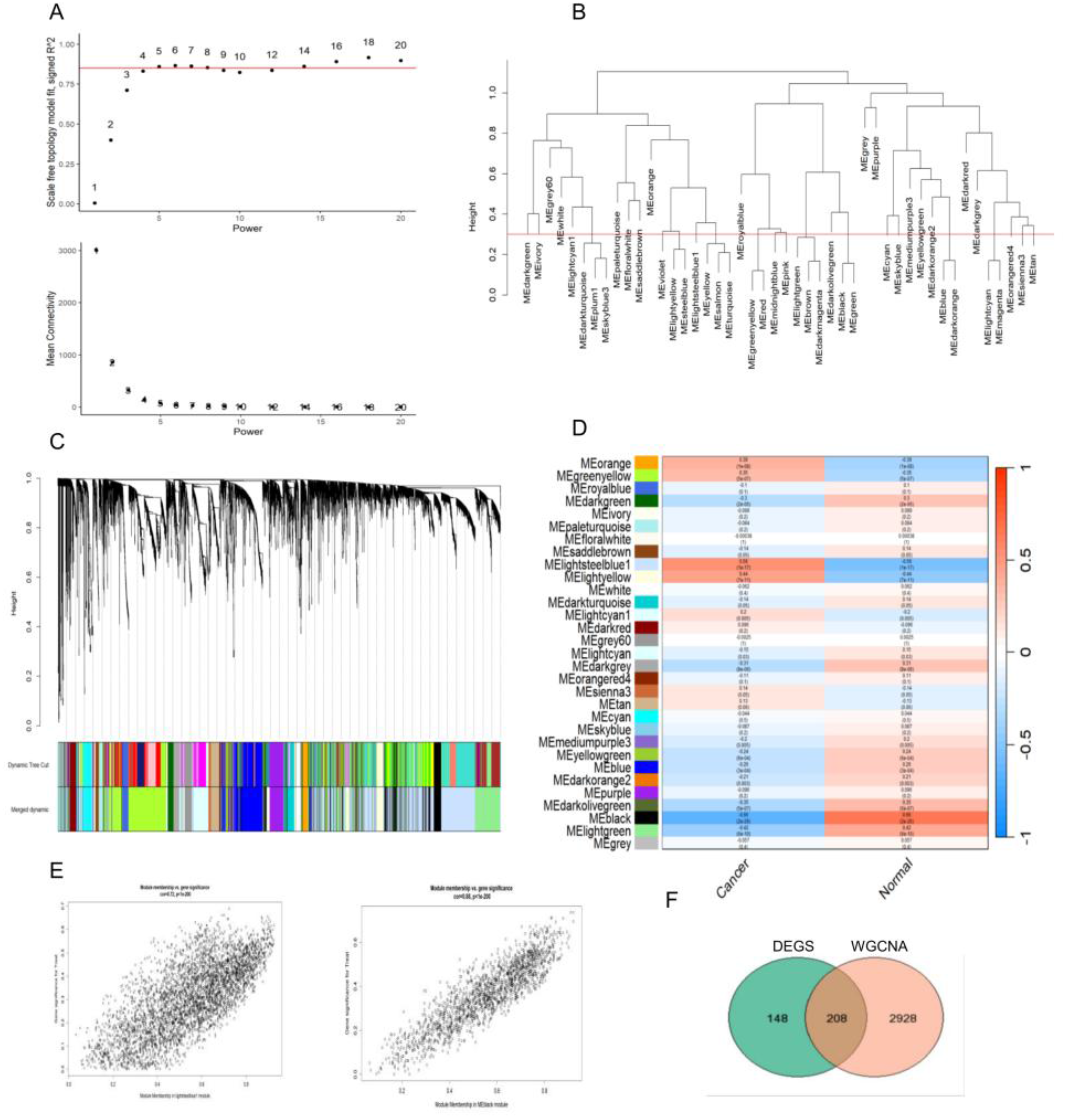
WGCNA-driven analysis of gene co-expression and module detection. (A) Scale-free fit index plot between power vs. signed R2 and mean connectivity plot. (B) Clustering of module eigengenes. (C) Dendrogram representing hierarchical clustering of the co-expressed gene set. (D) Heatmap of module eigengenes (rows) and clinical traits (columns) correlation. Each cell shows the correlation coefficient along with the corresponding p-value. (E) Module membership for lightsteelblue1 and black modules. (F) Overlap of DEGs with WGCNA module genes

Based on module eigengene correlation (threshold = 0.3) in Figure 3(B), similar expression profile modules were merged, resulting in a total of 31 modules. The cluster dendrogram of co-expressed genes is shown in Figure 3(C). Subsequently, modules were correlated with clinical phenotypes (cancer and normal) (Figure (3D)). The black and lightsteelblue1 modules show the highest correlation with TNBC status. The black module has a ModuleTraitCor of 0.66 and a ModuleTraitPvalue of 2E-26, while the light-steelblue1 module has a ModuleTraitCor of 0.56 and a ModuleTraitPvalue of 1E-17, respectively (Figure 3(E)). Finally, the genes from the black and lightsteelblue1 modules were intersected with the DEGs, resulting in 208 overlapping genes (Figure 3(F)).

### 3.3 GO term annotation and KEGG pathway characterisation

Enrichment analyses using GO and KEGG pathways were conducted to determine the biological roles of 208 potential genes that were identified by DEGs intersecting with two WGCNA modules (Figure 4(A) and (B)). The GO term BP included the mitotic cell cycle process, cell division, chromosome segregation, nuclear division, and sister chromatid segregation. Genes within the CC group were mainly enriched in structural components involved in mitosis, such as the spindle, mitotic spindle, chromosomal region, kinetochore, centrosome, and microtubule cytoskeleton. Additionally, the genes in the MF group show significant enrichment functions, microtubule binding, tubulin binding, microtubule motor-activity, ATP binding, and cyclin-dependent kinase regulator activity. KEGG pathway analysis identified key pathways, including cell cycle, PI3K-Akt signalling, and cellular senescence. Additional pathways included progesterone-mediated oocyte maturation, oocyte meiosis, ECM receptor-interaction, focal adhesion, homologous recombination, and the Fanconi anaemia pathway.

**Figure 4.**
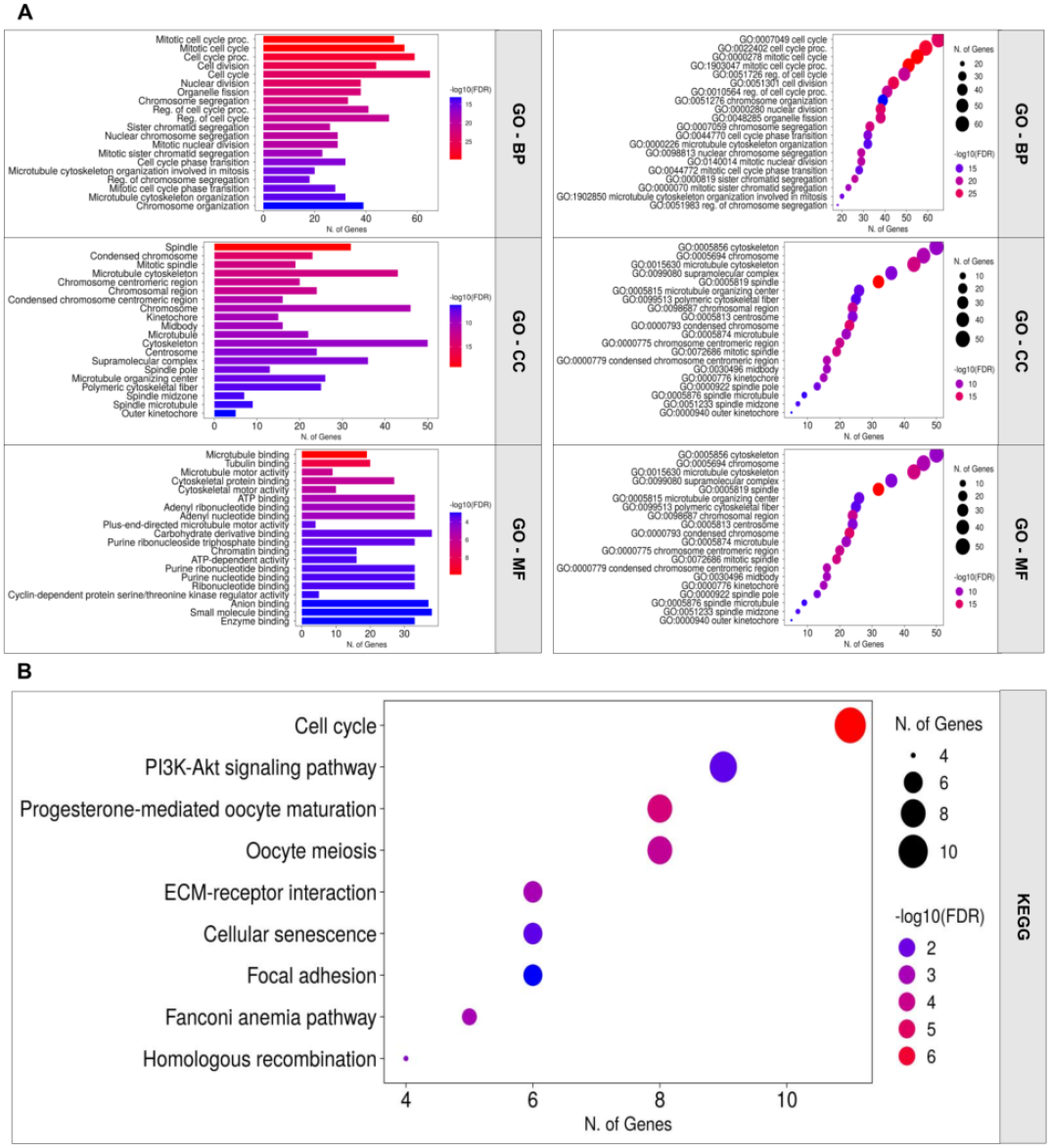
Functional enrichment analysis of 208 potential genes. (A) GO terms: categorised into BP, CC, and MF. (B) KEGG pathways showing enriched biological processes

### 3.4 ub gene identification via PPI network analysis

The PPI networks of black and lightsteelblue1 module genes mapped to DEGs were visualised using the Cytoscape CytoHubba plugin. Both modules’ top ten hub genes selected by the degree method are shown in Figure 5(A) and (B). The top five genes from each module (ten in total) were chosen for screening biomarkers for TNBC. These include cyclin A2 (*CCNA2*), kinesin family member 11 (*KIF11*), DNA topoisomerase II alpha (*TOP2A*), cyclin dependent kinase 1 (*CDK1*), cyclin B1 (*CCNB1*), matrix metallopeptidase 12 (*MMP12*), C-X-C motif chemokine ligand 8 (*CXCL8*), secreted phosphoprotein 1 (*SPP1*), neurotrophic receptor tyrosine kinase 2 (*NTRK2*), and myosin heavy chain 11 (*MYH11*). The combined PPI network of both modules is shown in Figure 5(C).

**Figure 5.**
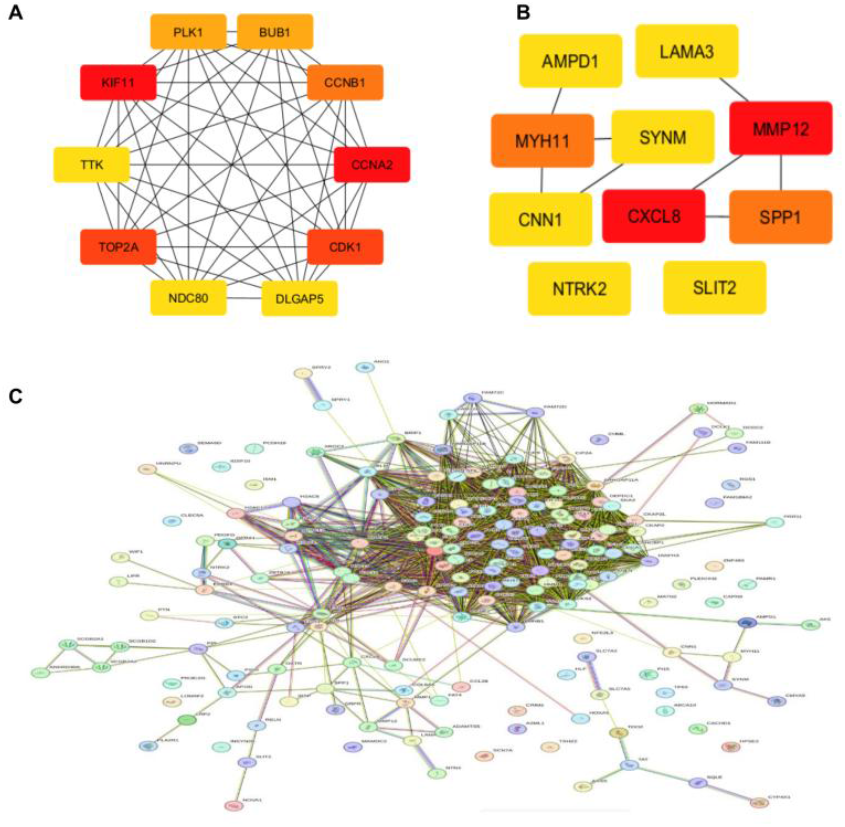
The top 10 hub genes using PPI interaction network analysis. (A) Black module (B) lightsteelblue1 module (C) combined PPI network

### 3.5 Machine learning-based significant gene selection

To investigate prospective biomarkers of TNBC, two machine learning feature selection models were applied to 208 critical genes based on cancerous and normal tissue samples. LASSO regression analysis determined the optimal value of λ = 13.878288, yielding 13 potential genes (Figure 6(A) and (B)). On the other hand, from SVM-RFE, the top ten significant genes were chosen from the output set of the SVM-RFE process (Figure 6(C) and (D)). Ultimately, intersecting the results from both ML approaches, three genes were consistently highlighted, including (*AASS*), cytoskeleton-associated protein 2 (*CKAP2*), and hepatic leukaemia factor (*HLF*) as prospective genes for screening biomarkers associated with TNBC (Figure 6(E)). These three genes were further validated by the ROC curve analysis; the AUC achieved were >0.94, proving their strong predictive value in distinguishing TNBC from normal tissues (Figure 6(F)).

**Figure 6.**
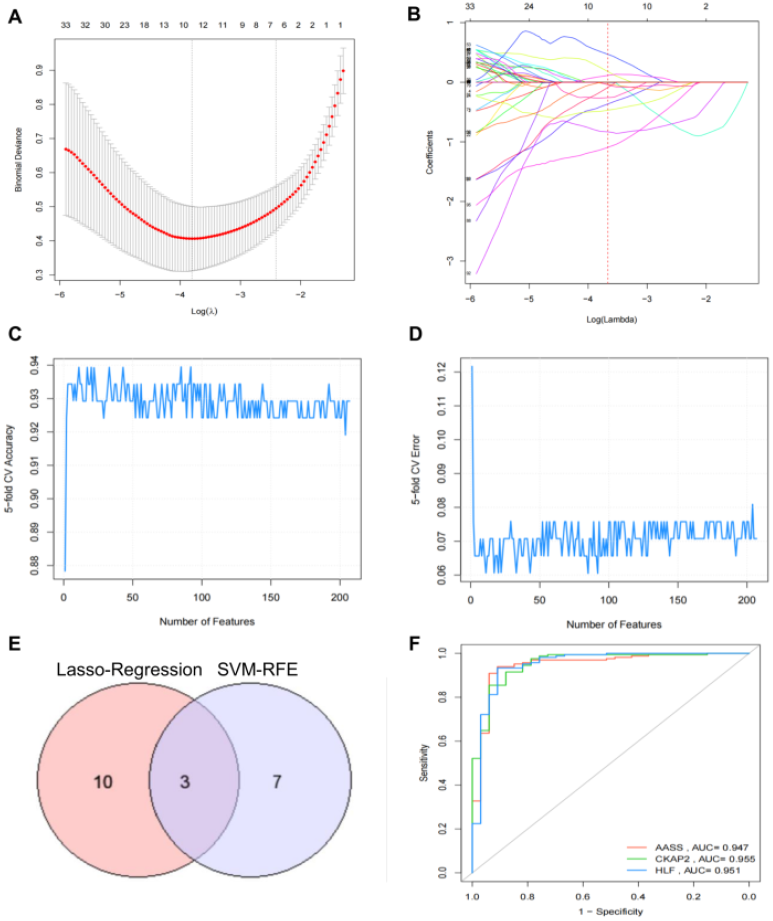
Identification and validation of biomarkers using LASSO regression and SVM-RFE. (A) Coefficient distribution of log-transformed λ values of the LASSO regression model. (B) Identification of significant genes with non-zero coefficients. (C) SVM-REF accuracy plot between 5-fold CV and number of features. (D) The SVM-RFE plot indicates the curves’ best accuracy and lowest error. (E) Intersecting biomarkers identified by ML (LASSO & SVM-RFE). (F) The ROC curve validates the predictive performance of the final selected biomarkers

### 3.6 Survival analysis on TNBC patient cohorts of prospective biomarkers

We used the KM plotter to validate the prognostic value of 10 hub genes and 3 key genes. This includes 534 TNBC patients for RFS analysis, 424 TNBC patients for DMFS, and 404 basal-type patients for OS analysis. Hazard ratios (HRs) < 1 indicated favourable outcomes, while HRs > 1 indicated unfavourable associations.

Among the favorable prognostic markers, *AASS* and *CCNA2* were consistently linked to improved survival outcomes across all analyzed endpoints, including RFS (HR = 0.66 and 0.62; P = 0.0061 and 0.011), OS (HR = 0.68 and 0.51; P = 0.047 and 0.00042), and DMFS (HR = 0.59 and 0.64; P = 0.012 and 0.022), respectively. Both genes demonstrated a protective effect in TNBC patients. In contrast, *CXCL8, SPP1*, and *CCNB1* were identified as unfavourable prognostic indicators. *CXCL8* was linked with a significantly high risk of poor outcomes across all survival metrics (RFS: HR = 1.79, P = 0.00024; OS: HR = 2.05, P = 0.00016; DMFS: HR = 1.62, P = 0.0073), while *SPP1* and *CCNB1* were also significantly correlated with adverse prognoses in RFS (HR = 1.39 and 1.9; P = 0.034 and 0.014) and DMFS (HR = 1.84 and 1.97; P= 0.00071 and 0.02), respectively. (Figure 7-9). Other genes showed partial or context-dependent prognostic value. For instance, *CKAP2* was significantly associated with improved OS, while *MYH11* showed contrasting effects associated with worse OS but better DMFS. Similarly, *CDK1* was protective in OS but showed borderline risk in DMFS. *KIF11* and *CDK1* were significantly associated with better OS, although their effects on other outcomes were weaker. These findings highlight five significant genes, including *AASS* and *CCNA2* as favourable and *CXCL8, SPP1*, and *CCNB1* as unfavourable prognostic indicators that may serve as potential biomarkers in TNBC. Their expression patterns are strongly linked to metastasis, disease recurrence, and overall survival outcomes of patients.

**Figure 7.**
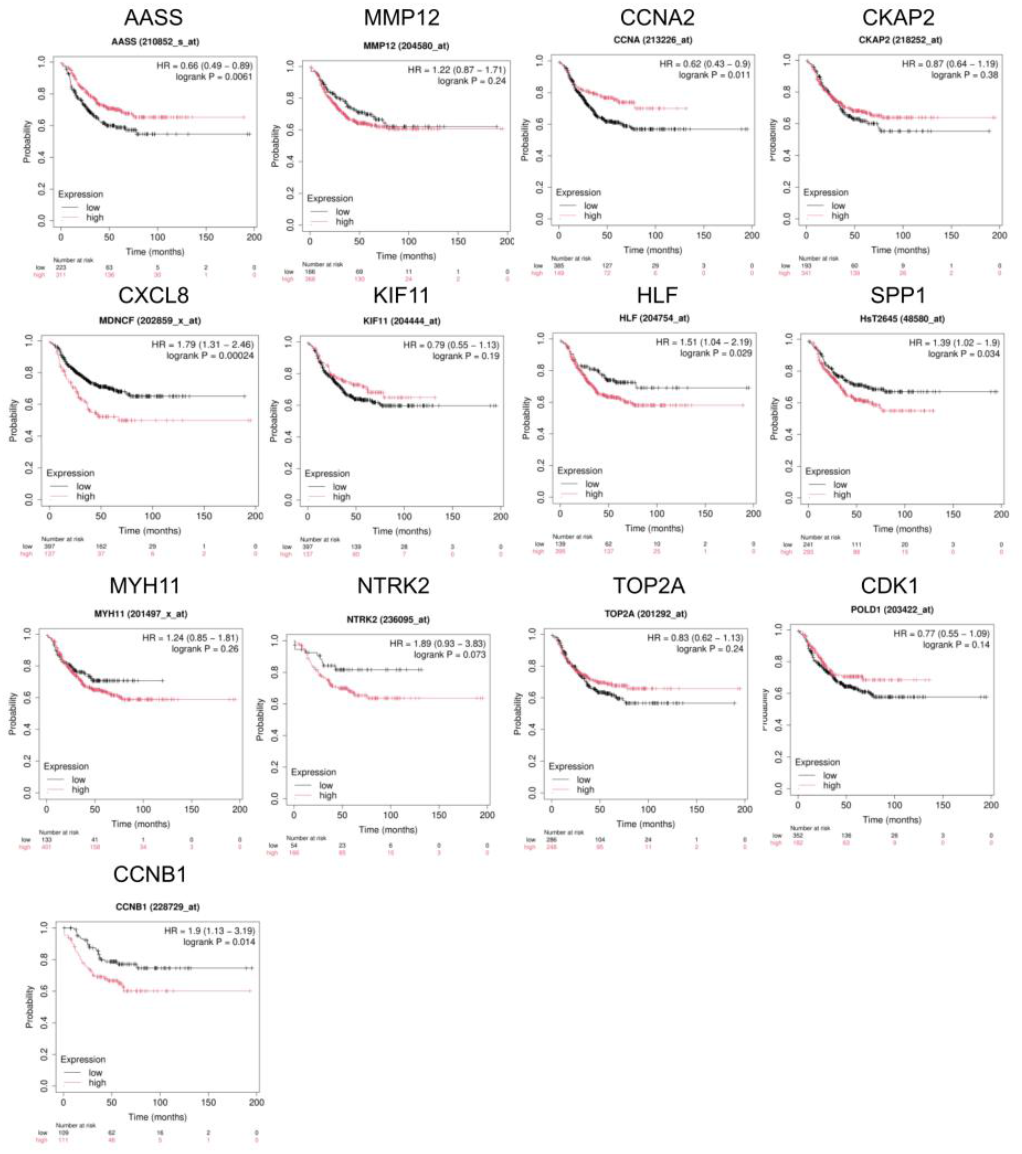
KM survival analysis of relapse-free survival (RFS) for candidate genes identified through PPI network topology and machine learning feature selection. Survival curves compare high- and low-expression groups stratified by median gene expression levels. Statistical significance was assessed using the log-rank test, showing the prognostic relevance of selected candidates in predicting TNBC recurrence risk.

**Figure 8.**
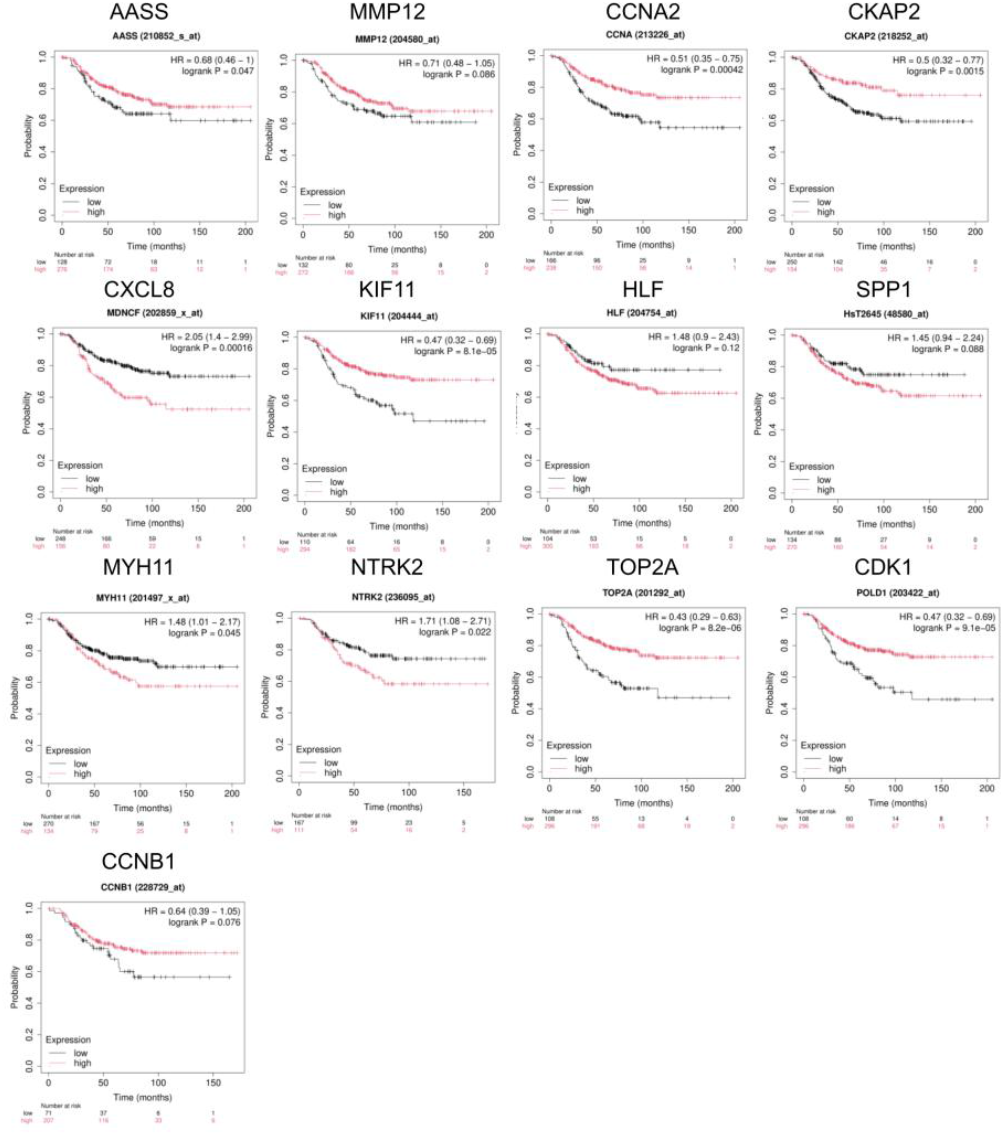
Overall survival (OS) analysis of hub and key genes identified, where patients were dichotomised into high- and low-expression cohorts, and survival differences were evaluated using the log-rank test, highlighting genes associated with favourable or poor overall prognosis in TNBC.

**Figure 9.**
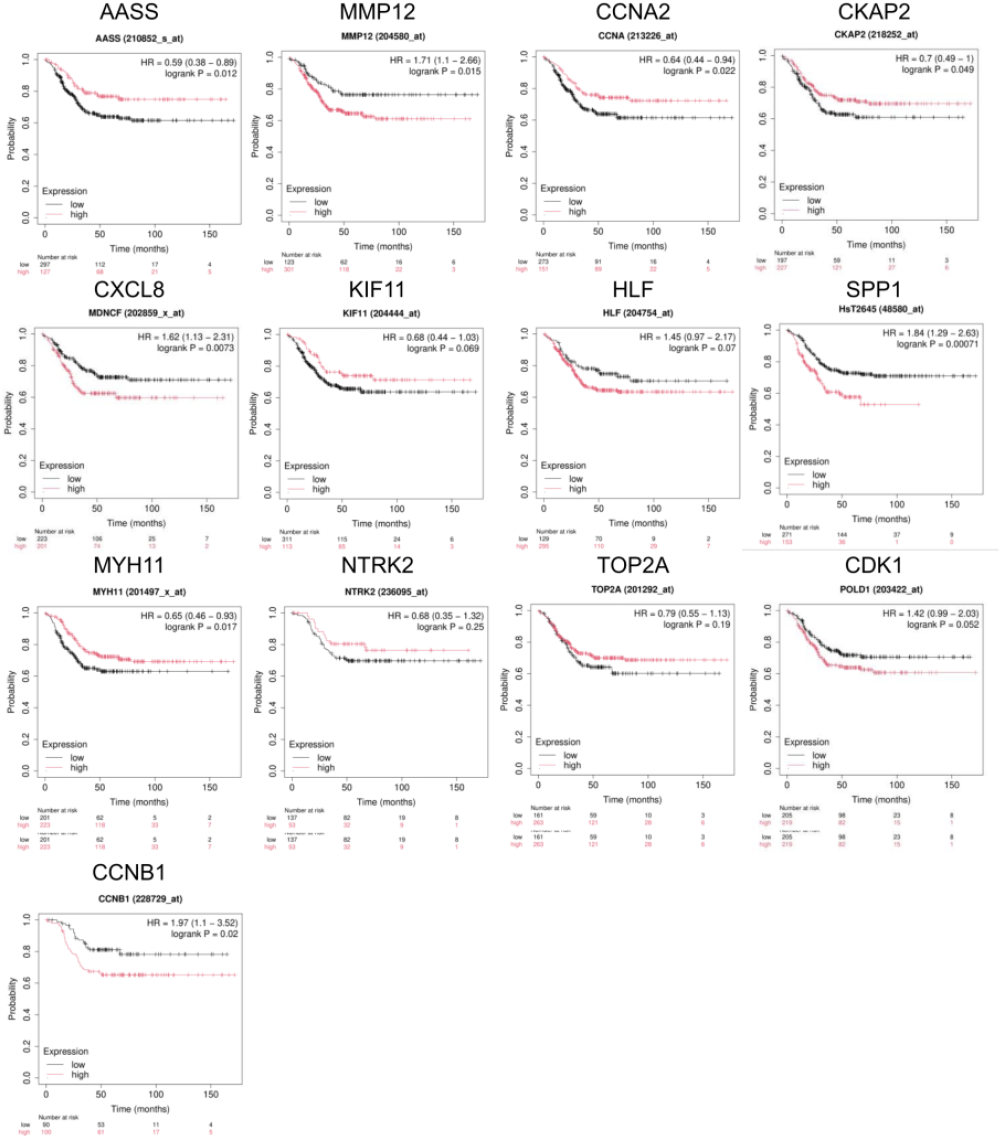
KM distant metastasis-free survival (DMFS) curves for biomarkers selected through combined PPI network modelling and ML-based feature selection. Expression-based stratification shows the statistically significant association between candidate gene expression and metastatic risk of progression.

### 3.7 Validation using independent TNBC cohorts

To strengthen the validity of findings, this study analysed DEGs from two additional independent external TNBC datasets (GSE38959 and GSE65194). A comparative analysis using Venn diagrams (Figure 10) revealed a shared subset of DEGs across all three datasets. Notably, the candidate genes *AASS, CCNA2, CXCL8, SPP1*, and *CCNB1* were consistently identified in the intersection, highlighting their potential significance and the reliability of our results.

**Figure 10.**
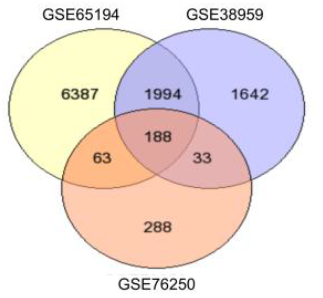
Venn diagram representing the overlapping DEGs identified between three independent TNBC datasets

### 3.8 AASS novel candidate in TNBC

The TNMplot gene chip and RNA-seq analysis revealed that *AASS* was significantly downregulated in BRCA compared to normal tissues. Further tumour and normal sample, stage-specific expression and subtype analysis confirm its consistent downregulation and tumour suppressor role, especially in TNBC Figure 11(A-H). UALCAN CPTAC also confirmed lower *AASS* protein expression across the BRCA subtype, Figure 11I. Immune Subtype Profiling demonstrated differential expression across six immune subtypes (C1– C6) and molecular subtypes, suggesting immune context-dependent roles. Figure 12a-d. Moreover, the Human Protein Atlas (HPA) indicates that *AASS* exhibits a protective role in other cancers Figure 11(J-L).

**Figure 11.**
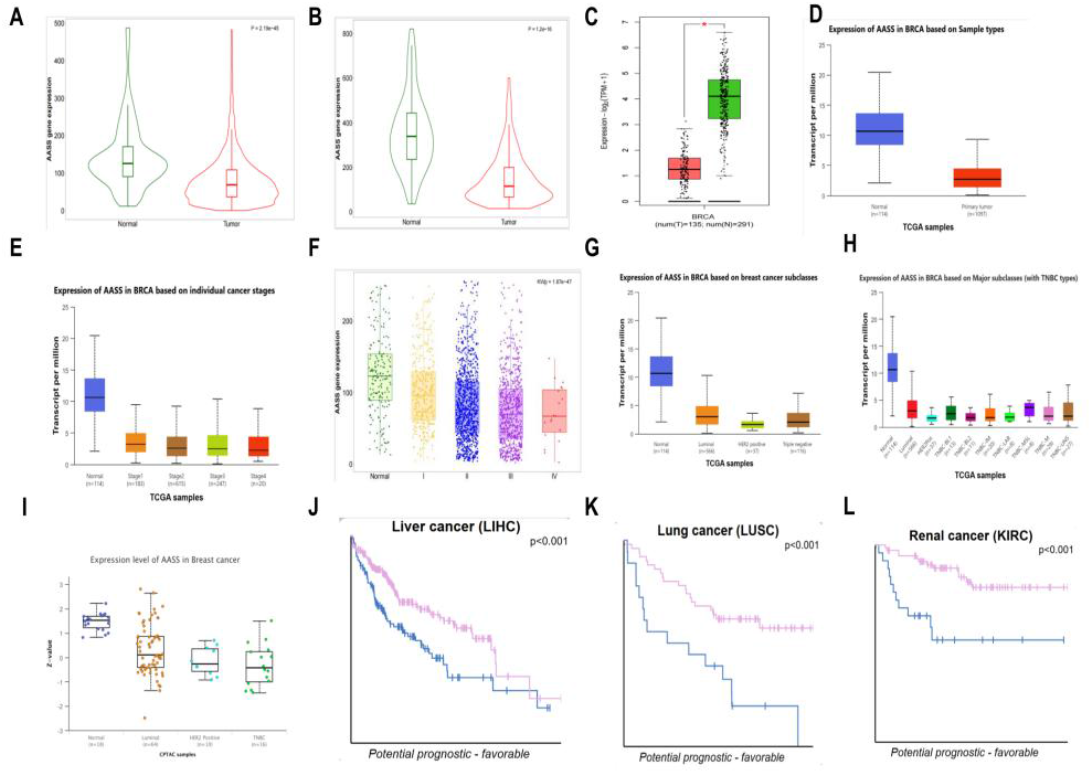
Extensive validation of *AASS*. Expression difference of *AASS* at the transcription level (A) Gene chip non-paired tumour and normal samples BRCA. (B) RNA seq Paired vs adjacent normal tissue samples BRCA. (C) Tumour vs. Normal BRCA by GEPIA2. (D) Normal vs Primary tumour BRCA UALCAN. (E) BRCA stage-specific analysis. (F) Stage-specific via TNMplot. (G) BRCA subtype analysis. (H) TNBC subtypes analysis. Expression difference at the protein level (I) CPTAC samples expression. Human protein atlas survival analysis (J) KM-Plot Liver cancer (K) KM-Plot Lung cancer. (L) KM-Plot Renal Cancer

**Figure 12.**
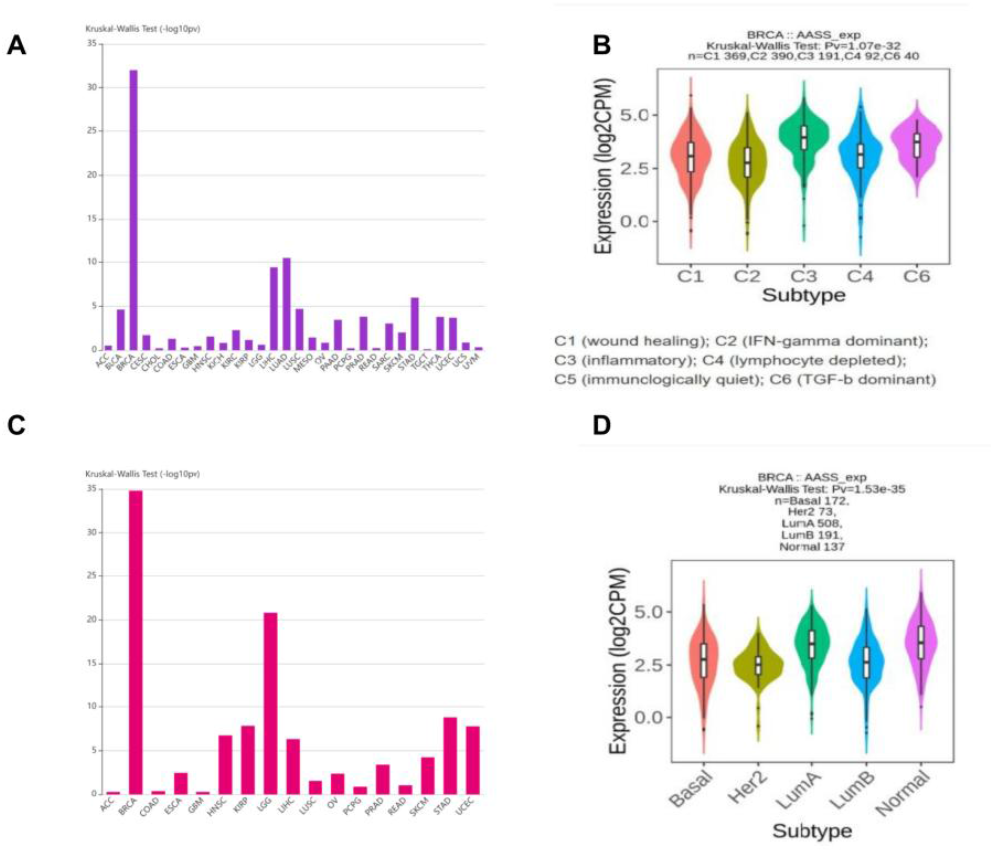
Immune and tumour microenvironment analysis (A) Immune subtypes and *AASS* expression association across cancers. (B) Immune subtypes and *AASS* expression are associated with BRCA. (C) Immune subtypes and *AASS* expression association across cancers. (D) Immune subtypes and *AASS* expression association across BRCA subtypes

## 4. Discussion

In this study, we first intersected DEGs with WGCNA lightsteelblue1 and black module genes and obtained 208 candidate genes associated with TNBC and normal samples in the GSE76250 dataset. GO and KEGG analyses showed that these genes were highly enriched in cell cycle regulation, mitotic spindle organisation, and microtubule-associated functions. Key enriched pathways included the cell cycle, PI3K-Akt signalling, and cellular senescence, along with additional roles in oocyte maturation, DNA repair, and extracellular matrix interactions. Moreover, from PPI networks based on degree scores calculated utilising the Cyto-Hubba module of Cytoscape, the ten most significant hub genes related to TNBC were selected, including *CCNA2, KIF11, CDK1, TOP2A, CCNB1, MMP12, CXCL8, SPP1, MYH11*, and *NTRK2*. Further analysis using ML techniques, including LASSO regression and SVM-RFE, was conducted on 208 candidate genes. As a result, three key genes were identified (*AASS, HLF*, and *CKAP2*). Subsequently, the prognostic potential of the top ten-ranked hub genes and three significant genes was further validated through analysis with KM-plotter.

In the KM Plotter, patients for RFS and DMFS were chosen based on negative ER and PR status determined by IHC and HER2 negativity confirmed by array data, ensuring the inclusion of pure TNBC cases. However, due to limited sample availability for OS analysis, the selection criteria were broadened to include patients classified as the St. Gallen basal-like subtype, which significantly overlaps with TNBC and is often used as a surrogate in molecular subtyping. Among the prognostic markers identified, *AASS* and *CCNA2* were consistently associated with improved outcomes across all survival endpoints (RFS, DMFS, and OS), suggesting their potential as favourable prognostic indicators. In contrast, *CXCL8, SPP1*, and *CCNB1* were identified as unfavourable prognostic genes. CXCL8 exhibited a markedly increased risk across RFS, DMFS, and OS; *SPP1* and *CCNB1* were significantly associated with poorer outcomes in DMFS and RFS.

The chemokine *CXCL8* is involved in the development and progression of different cancers, including breast, lungs, prostate, colorectal, and melanoma, due to its role in regulating tumour proliferation, invasion, and migration through paracrine or autocrine signalling mechanisms (Liu et al., 2016; Shen et al., 2021). High levels of *CXCL8* serum were reported to be linked with advanced disease stages, distant tumour metastasis, and larger tumours (Benoy et al., 2004). Conversely, during chemotherapy, low levels of *CXCL8* serum during chemotherapy have been linked with improved longevity, mainly in metastatic breast cancer (Tiainen et al., 2019). Research using breast cancer cell lines and xenograft models shows that *CXCL8* could enhance migration, invasiveness, and epithelial-mesenchymal transition in breast cancer cell lines (Nie et al., 2021). In our study of TNBC, *CXCL8* is frequently overexpressed and strongly associated with poor prognosis, promotes immune evasion and is resistant to chemotherapy. This would be worth investigating further.

*SPP1* is a multifunctional cytokine involved in cell adhesion, invasion, and immune cell recruitment, reported to be overexpressed in distinct tumours, including lung metastatic colorectal cancer and colorectal cancer associated with infliximab-resistant ulcerative colitis (Dai et al., 2023; Giannos et al., 2022), head and neck carcinoma, and gastric cancer, contributing to tumour progression, immune evasion, and chemoresistance (Feng et al., 2022; Lu et al., 2021). It is essential for cell invasion, migration, and proliferation; moreover, it modulates the tumour microenvironment and influences immune cell infiltration. In this study, high *SPP1* expression was associated with worse survival in TNBC, indicating its role as a mediator of metastasis and immune modulation. Further studies targeting *SPP1* or its downstream signalling could provide therapeutic benefits in TNBC.

*CCNB1* is a crucial molecule involved in regulating the G2/M stage in the cell cycle (Xia et al., 2021). Due to its role in mitosis, it is essential for tumour growth. *CCNB1* overregulation has been linked to poor patient survival outcomes in various cancer studies, including neutrophils in lung cancer and prostate cancer (Chen et al., 2022; Tang et al., 2023). In breast cancer, its expression level has been evaluated (Fu et al., 2022), and the association of *CCNB1* with the lack of hormonal receptors and the presence of HER2 receptors have been reported (Aljohani et al., 2022). Additionally, *CCNB1* involvement in TNBC has been discussed. In TNBC patients, compared to normal, over-expression of *CCNB1* is an unfavourable prognostic factor (Li et al., 2018; Lin, Wang and Yang, 2024). In our study, *CCNB1* overexpression was significantly associated with unfavourable outcomes, particularly in RFS and DMFS.

*CCNA2* (Cyclin A2) is part of a conserved cyclin family that controls the transition between G1/S and the G2/M stages of the cell cycle. It functions by making specific serine/threonine protein kinase holoenzyme complexes with cyclin-dependent kinases *CDK1* or *CDK2*. The cyclin component determines the substrate dependence of these complexes and selectively activates *CDK1* and *CDK2* at various phases of the cell cycle. It has prognostic significance in different cancers, including breast and colorectal cancer, as discussed in previous studies (Gan et al., 2018; Wang et al., 2021). *CCNA2* promotes the production of TNBC cells and serves as a potential biomarker for TNBC therapy (Lu et al., 2022). In this research, high expression of *CCNA2* was linked with improved survival rates in patients with RFS, DMFS, and OS, which calls for further in-depth research.

*AASS* plays an important role in the lysine degradation pathway, specifically converting lysine to acetyl-CoA via saccharopine. *AASS* in combination with *C1QTNF2* has recently been reported as a potential discriminative biomarker through integrative transcriptomic and machine learning approaches in uterine cancer (Golbaghi et al., 2024). Recent studies highlight its metabolic tumour-suppressive function. Activation of the lysine degradation pathway through overexpression of *AASS* and related enzymes significantly impaired proliferation in multiple cancer cell lines, indicating that these enzymes can restrain tumour growth by controlling metabolic flux through lysine catabolism (Dai et al., 2019). Complementing this, another study demonstrated that in human breast cancer cell lines, *AASS* overexpression led to a 5-fold increase in acetoacetate, a ketone body, which in turn induced autophagy and senescence, resulting in an 85% reduction in proliferation. Accumulation of acetoacetate appears to disrupt energy homeostasis in cancer cells, promoting cellular starvation and inhibiting growth. In breast cancer, the nuclear receptor PPARγ represses the expression of *AASS*, a key enzyme in lysine catabolism, by recruiting co-repressors such as *NCOR1, SMRT*, and *HDAC3* to the *AASS* promoter (Wu, Kramer and Crowe, 2023).

GO annotations suggests its localisation in key metabolic and regulatory compartments such as the mitochondria, cytosol, and nucleus, supporting its dual role in cellular metabolism and transcriptional control. Furthermore, functionally, *AASS* is not only involved in lysine catabolism but is also annotated with transcription co-repressor activity and negatively regulates transcription by RNA polymerase II, according to UniProt.

This dual functionality suggests that *AASS* may act at the interface of metabolic and epigenetic regulation. By contributing to transcriptional repression, *AASS* may inhibit genes that function in cell cycle progression, angiogenesis, or immune evasion. Simultaneously, its role in the lysine degradation pathway may regulate cellular energy balance and redox status, further supporting its function as a metabolic gatekeeper in tumour biology. This regulatory function is particularly relevant in TNBC, where receptor deficiency leaves tumours highly reliant on alternative oncogenic pathways. By negatively regulating gene expression, *AASS* might suppress the transcription of oncogenes or genes supporting metabolic reprogramming, thereby acting as a tumour inhibitor at the transcriptional level.

Together, these observations and literature support highlighting *AASS* as a best-fit candidate tumour suppressor with both metabolic and gene regulatory functions. At present, *AASS* has not been previously reported as a biomarker in TNBC. Our findings are the first to highlight its potential involvement in TNBC, where it may serve as a tumour inhibitor. In summary, our results mention that *AASS* and *CCNA2* are protective prognostic indicators in TNBC, whereas *CXCL8, SPP1*, and *CCNB1* are linked to worse clinical outcomes that may act as potential biomarkers for early detection and therapeutic targeting in TNBC.

## 5. Conclusion

In this study, an integrated approach combining bioinformatics expression analysis with ML feature selection techniques was utilised to identify potential biomarkers in TNBC. Identified genes showed strong involvement in cell cycle regulation and cancer-related pathways through functional enrichment analysis. PPI network analysis and ML identified several key genes, which are verified by two independent datasets, including *AASS, CCNA2, CXCL8, SPP1*, and *CCNB1*. Survival analysis showed *AASS* and *CCNA2* as favourable, while *CXCL8, SPP1* and *CCNB1* were associated with poor outcomes. *AASS* emerged as a novel candidate. Further extensive multilevel validation positioned it as a novel biomarker for TNBC. Despite this finding, future experimental investigation of *AASS* is crucial to elucidate the molecular mechanisms underlying TNBC, which may provide valuable insights into TNBC biology.

## Data availability

Data utilised and analysed during the study are openly available in Gene Expression Omnibus (GEO, https://www.ncbi.nlm.nih.gov/geo/).

